# Characterization of rumen microbiome and metabolome from an oro-esophageal probe and fluid, particulate and fluid-particulate fractions from rumen fistula in Holstein dairy cows

**DOI:** 10.1101/2022.07.25.501495

**Authors:** Lais L. da Cunha, Hugo F. Monteiro, Igor F. Canisso, Rodrigo C. Bicalho, Felipe C. Cardoso, Bart C. Weimer, Fabio S. Lima

**Author notes:** Corresponding author: Fabio S. Lima, University of California, Davis, CA, 95616, USA.

## Abstract

Less invasive rumen sampling methods such as oro-esophageal probes became widely popular to explore the rumen microbiome and metabolome. However, it remains unclear if such methods represent well the rumen contents from fluid and particulate fractions. Herein, we characterized the microbiome and metabolome in rumen content collected by an oro-esophageal probe and fluid, particulate, and the combined fluid-particulate fractions collected by rumen fistula in ten multiparous Holstein dairy cows. The 16S rRNA gene was amplified and sequenced using the Illumina MiSeq platform. Untargeted metabolome was characterized using gas chromatography of a time-of-flight mass spectrometer. Although the pH of oro-esophageal samples was greater than those of fluid, fluid-particulate, and particulate ones, we found no difference in alpha and beta-diversity of their microbiomes. Bacteroidetes, Firmicutes, and Proteobacteria were consistently the top three most abundant phyla representing ~90% of all detected phyla across all samples. The overall metabolome PLS-DA of oro-esophageal samples was similar to the fluid-particulate samples but differed from fluid and particulate. Enrichment analysis pathways revealed few differences between oro-esophageal and fluid-particulate samples, such as the synthesis of unsaturated fatty acids. The results of the current study suggest that oro-esophageal sampling can be a proxy to screen the rumen microbiome with the 16S platform and overall fluid-particulate metabolome for a single-time and diet context. Nonetheless, studies focusing specifically on fluid and particulate metabolomes and specific metabolic pathways should carefully consider the sampling method used.

**IMPORTANCE:** The techniques used to collect the rumen contents (oro-esophageal probe and rumen fistula) suggested potential differences in the populations of rumen microbes, and the implications of these techniques for high throughput studies characterizing the rumen microbiome and metabolome need further elucidation. Ten rumen-fistulated Holstein dairy cows were used to characterize the microbiome and metabolome of samples collected using an oro-esophageal probe and the rumen-fistula fluid, particulate, and fluid-particulate fractions. The results of the current study suggest that oro-esophageal sampling represents well the rumen microbiome and overall fluid-particulate metabolome. However, fluid and particulate metabolomes and specific metabolic pathways across all types of rumen samples differed, indicating that studies focused on the characterization of rumen metabolome variable fractions should carefully consider the sampling method used.

## INTRODUCTION

The ruminant digestive tract is composed of four-chambered stomachs degrading and processing the diet ingested by animals (1). This process depends on predominantly anaerobic microorganisms inside the rumen responsible for breaking down a variety of feed particles into digestible nutrients, such as β-linked carbohydrates into digestible sugars (2). Furthermore, fermentation of these nutrients by ruminal microorganisms is advantageous to their growth and proliferation and provides significant precursors for the host’s metabolic pathways (3). The diet degradation process through fermentation by microorganisms starts in the particulate fraction of the rumen content. Microorganisms such as *Fibrobacter* and *Ruminococcus* adhere to fibrous polysaccharides and hydrolyze them through biofilm formation into di- and monosaccharides (2). In the fluid fraction, fermentation of smaller molecules intensifies, and end-products of fermentation are highly produced by microorganisms and used by the host in metabolic processes (4). Therefore, the adequate characterization of the ruminal microbiome is essential for developing efficient diagnostic tools and therapeutic interventions to improve animal health and advance the current knowledge on major nutritional issues faced by the dairy and beef industries (5, 6).

Historically, rumen fistula was the gold-standard method used to investigate the interactions of microbes and feed in these forestomaches that play a pivotal role in ruminants’ digestion (3). Recently, several studies used less invasive methods to characterize high-throughput data in large populations (7–9). However, it remains unclear how these less invasive methods represent microbes and metabolites associated with specific fractions of rumen content (i.e., particulate and fluid) previously reported in rumen-fistulated studies (10). Amongst the less invasive methods, a collection using an oro-esophageal probe (7) became a popular technique to obtain rumen contents without the need for major surgery and associated risks and cost with rumen fistulation. The advance in sequencing methods led to studies investigating various aspects of the rumen microbiome and metabolome (11–14). However, studies comparing the rumen microbiome derived from oro-esophageal and rumen fistula sampling techniques have been controversial suggesting either consistent (15, 16) or biased results for between these two techniques (17, 18). Furthermore, no study assessed the rumen metabolome concurrently with rumen microbiome for these different sampling techniques to assess the feasibility of the less invasive oro-esophageal approach for multi-omics studies.

Considering the necessity of a large number of animals for proper characterization of cows’ genotype and phenotype for rumen multi-omics studies, we propose a study to test the hypothesis that samples collected using an oro-esophageal probe yield a microbiome and metabolome similar to combining fluid and particulate fractions collected using a rumen fistula, but it is distinct from each fraction alone. The aims of the study were to characterize the microbiome and metabolome of rumen samples collected using an oro-esophageal device and rumen fistula particulate, fluid, and combined fluid-particulate fractions.

## MATERIALS AND METHODS

All experimental procedures were conducted following protocols approved by the University of Illinois at Urbana-Champaign Institutional Animal Care and Use Committee (Protocol 17172). Ten fistulated multiparous Holstein lactating cows averaging 688 ± 78 kg BW were enrolled in the study. All cows were housed in a tie-stall system with sand bedding, fed twice a day ad libitum, and had free water access at all times. Diets were formulated using AMTS.Cattle.Pro version 4.7 (2017, AMTS, LLC, Groton, NY) to meet or exceed recommendations for cows producing 41 kg of milk/d with a target of 3.8% milk fat and 3.2% milk protein and a predicted DMI of 25 kg/d. The diet fed consisted of corn silage, alfalfa hay, soybean meal, dry ground corn grain, canola meal, corn gluten feed, soy hulls, dried molasses, bypass fat, premixed minerals (dicalcium phosphate, calcium, potassium carbonate, sodium bicarbonate, potassium chloride, magnesium oxide), rumen-protected lysine and methionine, urea 46%, salt white, and vitamins.

### Sampling procedure

Rumen samples were collected 5-6 after morning feeding, and samples (oro-esophageal content, fluid, particulate, and combined fluid-particulate fistula) were collected from each of the ten cows enrolled in the study totalizing 40 samples. Briefly, an oro-esophageal sampling device was used to collect rumen content samples (7). A vacuum pump equipped with a glass container was connected to a probe of approximately 200 cm in length and 2.5 cm in diameter before being used. The probe was inserted orally in the cows until it could reach the rumen. Rumen content was collected through building vacuum pressure in the probe. The first two samples were discarded to avoid contamination of rumen contents with esophageal components, such as saliva and mucus. Then, approximately 500 mL of rumen content was collected, and 15 mL of the content was immediately stored in sterile conical tubes. Before the sample collection from each cow, the oro-esophageal tube was thoroughly cleaned to avoid cross-contamination. Samples from rumen fistula representing the combined cranial, caudal, dorsal, and ventral sacs were collected according to their respective fractions: fluid and particulate fractions separately and a homogeneous sample containing both fractions as a proxy to the overall composition of rumen contents. In brief, a homogenous fraction was collected and squeezed through two layers of cheesecloth saving the fluid (15 mL) and particulate (50 mL) contents in separated containers. Then, a homogenous fraction was collected through the rumen containing 50 mL of the combined fluid and particulate contents. During all collections, ruminal pH was measured using a portable pH meter immediately after sampling. Samples were immediately frozen and transported to the laboratory in Urbana, IL, where they were kept at −80°C freezer until further analyses.

### DNA extraction, library preparation, and sequencing

Bacterial DNA was extracted similarly to Lima et al. (2015). Briefly, rumen samples were thawed at 4°C and later centrifuged for 10 min at 16,000 RCF in a DNase-free microcentrifuge tube. The supernatant was discarded, and the pellet was resuspended in nuclease-free water. A QIAamp PowerFecal DNA Extraction Kit (Qiagen) was used for genomic DNA isolation. Except for the addition of 400 mg of lysozyme during bacterial resuspension and the following incubation of 12 h at 56°C to maximize bacterial DNA extraction, all other manufacturer’s instructions were followed for genomic vDNA isolation. A NanoDrop ND-1000 spectrophotometer (NanoDrop Technologies, Rockland, DE, USA) was used later at wavelengths 230, 260, and 280 nm for DNA concentration and purity measurements.

Library preparation and sequencing were performed similarly to those described by Kozich et al. (19). Amplification was performed through polymerase chain reaction (PCR) in a Bio-Rad C1000 TouchTM Thermal Cycler (BIO-RAD, Hercules, CA, USA). The V4 region of the 16S rDNA gene was amplified using dual-index (forward and reverse) bacterial primers through an initial 95°C denaturation for 5 min, followed by 30 cycles of 30 s at 95°C, 30 s at 55°C, 1 min at 72°C, and 5 min for final elongation at 72°C. Primers and small DNA fragments were removed using a 1% low melting agarose gel extraction kit (National Diagnostics, Atlanta, GA, USA). Purification and normalization of amplicons were performed using a SequalPrep plate kit (Invitrogen, USA), and the DNA concentration was measured with a Qubit^®^ Fluorometer. Adapters were added to the amplicons, and a DNA library was prepared by equally pooling them together; qualitative real-time PCR was used for a quality check. A total of 40 samples were sequenced using an Illumina MiSeq 2500 platform. Sequences were deposited in the Sequence Read Archive (SRA) of the National Center for Biotechnology Information (NBCI) under access number PRJNA784126.

### Metabolomics Data Acquisition and Processing

Ruminal metabolites were extracted following the procedure of Fiehn et al. (2008) and analyzed in gas chromatography of the time-of-flight mass spectrometer (GC-TOF) (20). The retention index and the complete mass spectrum were encoded as a string. All thresholds reflect settings for ChromaTOF v. 2.32. Quantification was reported as peak height using the unique ion as default unless a different quantification ion was manually set in the BinBase administration software BinView. We detected 185 known metabolites from a total of 421 untargeted primary metabolites found in our analysis. A column of 30 m length by 0.25 mm internal diameter with 0.25 μm film made of 95% dimethyl/5 diphenyl polysiloxanesne was used in a Restek corporation Rtx-5Sil MS. The gas helium (99.99% purity) was used a carrier for the analysis, and the column temperature was set between 50 – 330°C at flow-rate of 1 mL min^-1^. The oven temperature was set to 50°C for 1 min, then ramped at 2Ü°C min^-1^ to 330°C, and held constant for 5 min. Finally, the injection temperature was set to 50°C and ramped to 250°C by increments of 12°C^-1^. The retention of primary metabolites (amino acids, hydroxyl acids, carbohydrates, sugar acids, sterols, aromatics, nucleosides, amines, and miscellaneous compounds) were evaluated.

### Bioinformatics and statistical analyses

The first step in our bioinformatic analyses was the preparation of our metadata. For that, different sources of rumen samples were the groups for comparison. Downstream analysis was performed by testing differences between bacterial communities of each group created in the metadata. Upstream and downstream analyses of the sequenced amplicons were mostly performed in R Studio 2021.09.1. Sequences were denoised using the *dada2* pipeline (21), in which demultiplexed fastq files were inspected, filtered, and trimmed based on their quality scores and error rates. Chimeras were removed and an ASV table was created. Taxonomy was assigned using the 16S rRNA SILVA v138 database (22) with the *phyloseq* package (23). Total taxa were then split into taxonomy levels, and the relative abundance of the ASVs within each taxonomy level was calculated using the *phyloseq* package.

Alpha-diversity indexes [(total sequences, chimeras, unused sequences, Shannon, Chao 1, Inverse Simpson, and Rarity (low and rare abundant taxa)] were calculated using the *microbiome* and *vegan* packages (24, 25). Prevalence interval for microbiome evaluation [PIME; *pime* package (26)] was used to better select statistically and biologically relevant taxa for beta-diversity analysis. This latter pipeline allows the filtering of noise within each group by using random forest classification. Based on an appropriate prevalence interval calculated for the tested groups, taxa that were not shared within the same group were removed for a better visualization of the differences among bacterial communities.

Using the final PIME filtered taxa, a principal coordinate analysis (PCoA) was performed, and a graph was generated using the *ggplot2, dplyr, hrbrthemes, viridis, ggsci*, and *RColorBrewer* packages. Similarly, a permutational multivariate analysis of variance [PERMANOVA; (27)] was performed to test bacterial community’s dispersion difference. Linear discriminant analysis of effect size [LEFSe; (28)] was used to evaluate taxa differences between each sampling procedure. The LEFSe algorithm is based on three statistical tests (Kruskal-Wallis and Wilcoxon sum-rank tests, and linear discriminant analysis) to declare taxa differences in bacterial communities. However, no difference was observed in the LEFSe analysis and, thus, no results were reported in this study.

Metabolomic analyses were performed using Metaboanalyst 5.0 (www.metaboanalyst.ca). In brief, partial-least square discriminant analysis (PLS-DA), pathway, and enrichment analyses were performed to understand differences among compared groups. The KEGG metabolites library was used, and the top 5 enriched pathways were considered for direct comparisons between the oro-esophageal procedure (newer method of rumen sampling) and the fistula-fluid and particulate group (traditional gold standard for rumen sampling).

Lastly, a model containing the fixed effect of sampling procedure and the random effect of cow was fitted in SAS 9.4 for pH, all alpha-diversity variables, and the relative abundance of bacterial taxa. Statistical analyses were considered significant when *P* ≤ 0.05. When a statistical difference was observed, Tukey-Kramer test was used to compare group means, and the same *P*-value threshold was used to define in between-group differences.

## RESULTS

### Descriptive data sequencing results and rumen pH

There was no significant difference in the quality of total sequencing data analysis amongst sampling methods and fraction type. Dairy cows in the study had, on average, 39,292 unique sequences identified with no statistical significance (*P* = 0.44) amongst the four sampling groups compared (Fig. 1A). Likewise, no differences in sampling groups were present for the number of chimeras (*P* = 0.15, Fig. 1B) and the number of unused sequences (*P* = 0.67, Fig. 1C), indicating that these methodologies do not differ in the addition of noise to microbiome analysis. A total of 7,416 taxa were identified after taxonomy assignment, which was used for later downstream analyses.

**Figure 1.**
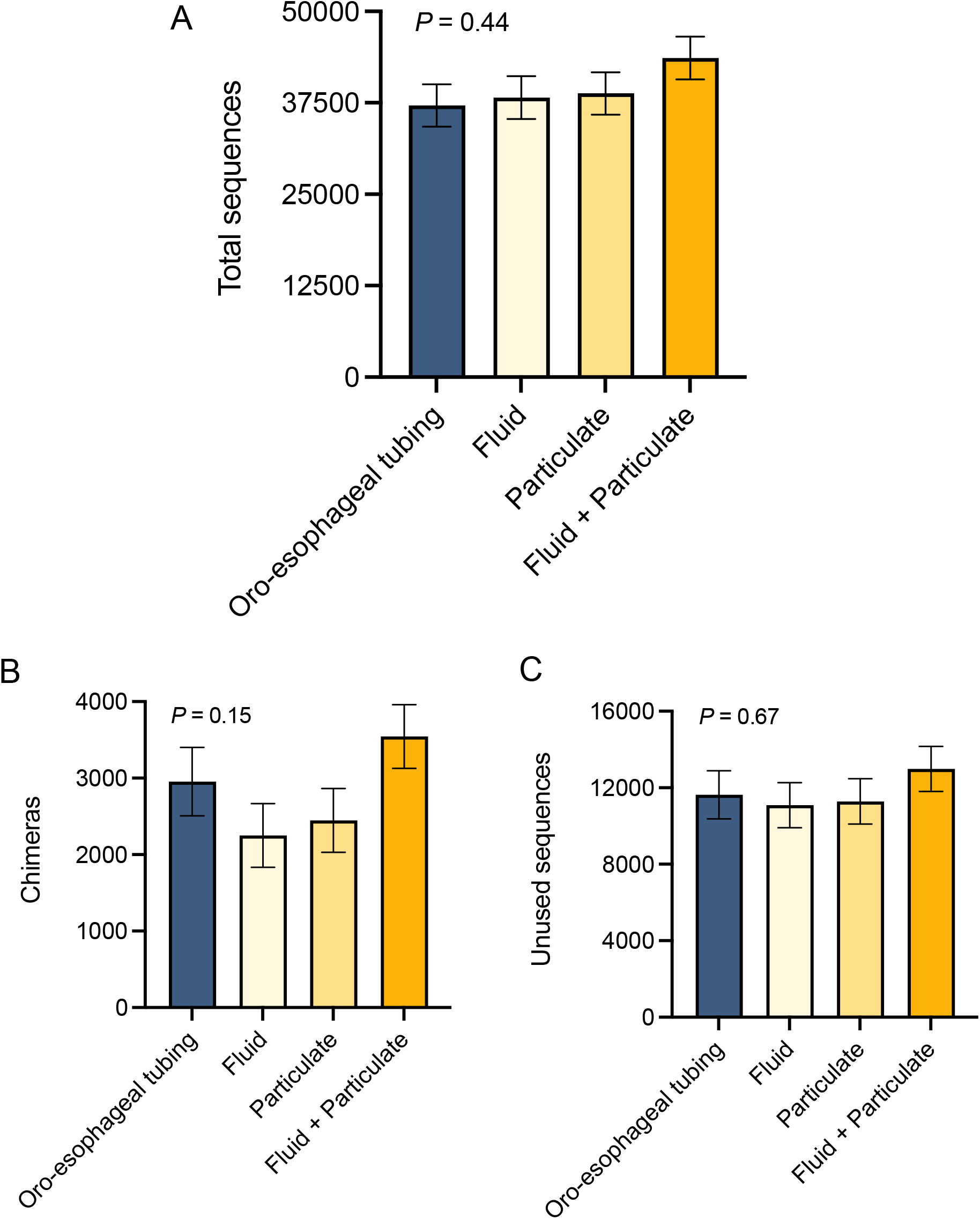
(A) total sequences (B) chimeras (C) unused sequences amongst fluid from oro-esophageal tubing and fluid, particulate and combined fluid-particulate collected from fistula. Statistical differences were declared at *P* ≤ 0.05. Different superscripts mean groups differ through Tukey-Kramer test performed at *P* ≤ 0.05 significance level.

The pH ranged from 6.3 to 6.8 amongst different sampling methods and fractions (fluid, particulate, and combined fluid-particulate), as shown in Fig. 2A. Overall, ruminal fluid pH collected through an oro-esophageal probe was greater (*P* < 0.01) than the pH of ruminal contents collected by rumen fistula (Fig. 2A).

**Figure 2.**
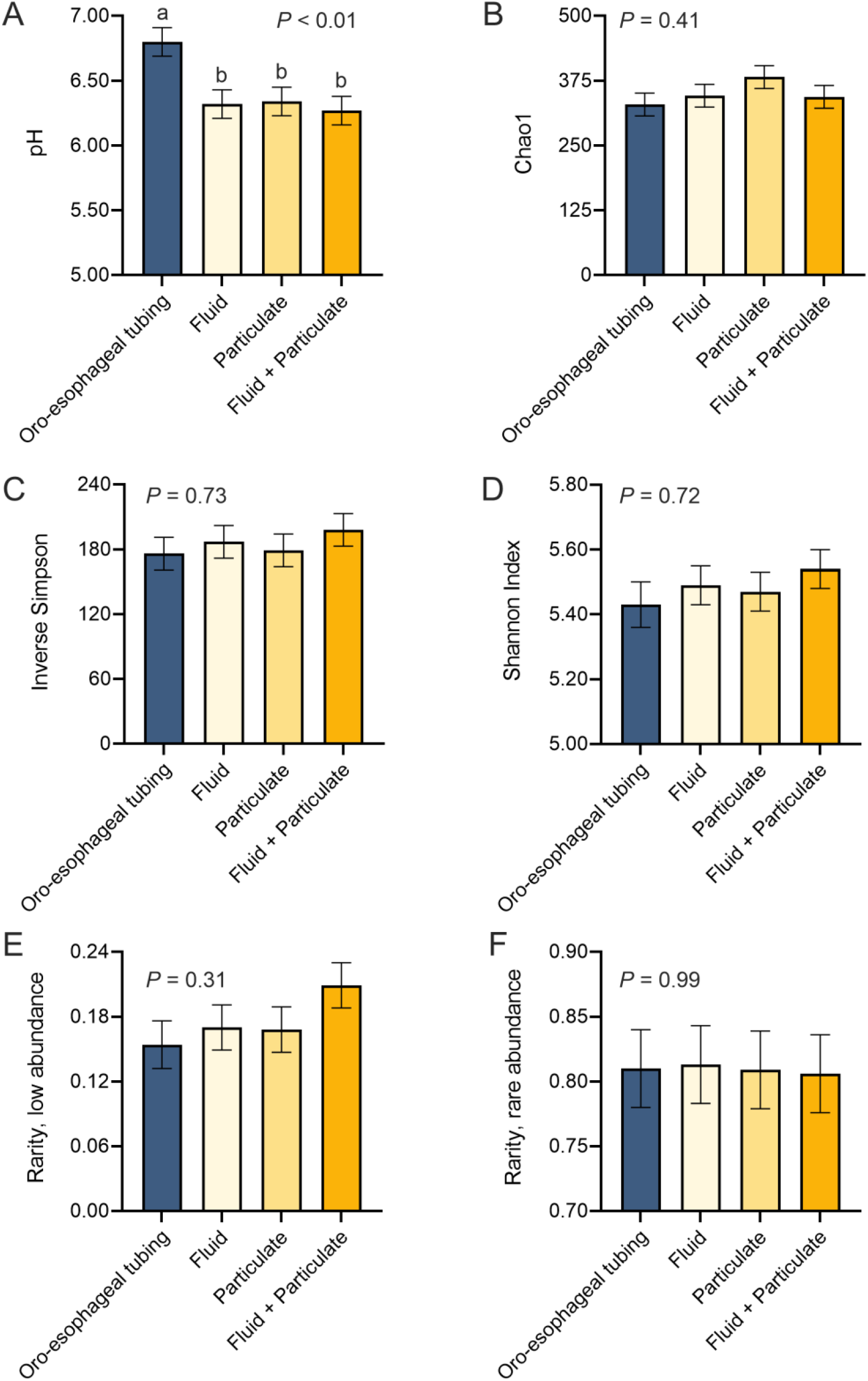
(A) pH (B) Chao1 (C) Inverse Simpson (D) Shannon Index (E) rarity, low abundance (F) rarity, rare abundance amongst fluid from oro-esophageal tubing and fluid, particulate and combined fluid-particulate collected from fistula. Statistical differences were declared at *P* ≤ 0.05. Different superscripts mean groups differ through Tukey-Kramer test performed at *P* ≤ 0.05 significance level.

### Rumen microbiome

No differences were detected in the rumen microbiome composition among the different sampling methods, as depicted in the results of the principal coordinate analysis (Fig. 3). Permutational multivariate analysis of variance and LEfSe also revealed no significant different taxa between oro-esophageal samples and rumen fistula fractions. Furthermore, no difference was present amongst sampling methods for the 14 identified phyla. The ten most abundant phyla mean relative abundance (MRA) in the rumen are reported in Table 1. The most common phyla across all samples were Bacteroidetes, followed by Firmicutes. No differences in the 30 most prevalent genera mean relative abundance was detected amongst sampling methods as well, nor were there differences amongst rumen source in the discriminant analysis. Prevotella, Bacteroides, Succiniclasticum, Treponema, Ruminococcus, and Butyrivibrio were the most prevalent genera.

**Figure 3.**
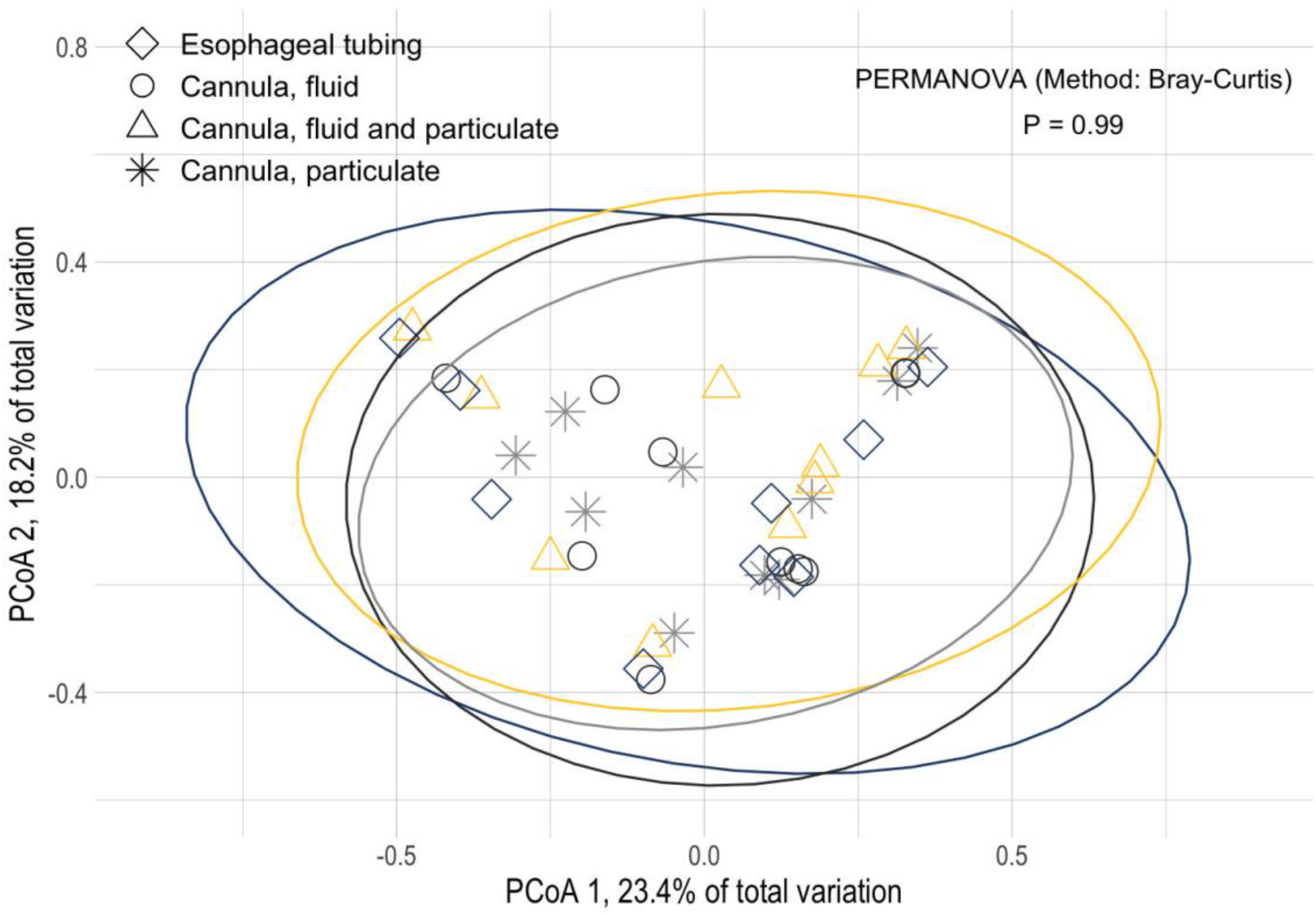
Principal coordinate analysis (PCoA) of the bacterial community composition using Bray-Curtis distance from ruminal microbial composition samples collected through an oro-esophageal tubing, and the fractions fluid, particulate, and combined fluid-particulate from rumen fistula. Statistical differences for permultational multivariate analysis of variance (PERMANOVA) was declared at *P* ≤ 0.05.

**Table 1.** Effects of different procedures to collect rumen fluid in lactating dairy cows on phylum relative abundance in the rumen.

| Phylum, % | Samples |  |  |  | SEM | *P*-values<sup>1</sup> |
| --- | --- | --- | --- | --- | --- | --- |
|  | Oro-esophageal | Fistula Fluid | Fistula Particulate | Fistula Fluid + Particulate |  |  |
| Bacteroidetes | 54.4 | 57.0 | 55.6 | 54.1 | 3.73 | 0.95 |
| Firmicutes | 29.8 | 29.8 | 29.0 | 32.1 | 5.19 | 0.97 |
| Proteobacteria | 6.62 | 5.65 | 6.50 | 6.24 | 2.90 | 0.99 |
| Spirochaetotes | 4.21 | 2.99 | 3.13 | 3.03 | 0.94 | 0.78 |
| Euyarchaeotes | 1.28 | 1.77 | 1.76 | 1.25 | 0.37 | 0.61 |
| Fibrobacterotes | 1.35 | 1.02 | 1.38 | 0.89 | 0.52 | 0.89 |
| Verrucomicrobiotes | 0.45 | 0.68 | 0.53 | 0.39 | 0.11 | 0.28 |
| Patescibacteria | 0.49 | 0.46 | 0.48 | 0.55 | 0.06 | 0.70 |
| Actinobacteriotes | 0.23 | 0.38 | 0.43 | 0.27 | 0.16 | 0.78 |
| Planctomycetes | 0.24 | 0.35 | 0.35 | 0.35 | 0.14 | 0.94 |

<sup>1</sup>Significant differences were considered at $P \leq 0.05$ , and a tendency was between $P > 0.05$ and $\leq 0.10$ .

The richness and diversity indexes are described in Fig. 2B–2F. The Chao1 richness, Shannon diversity index, Inverse Simpson, rarity, low, and rare abundance did not differ amongst oro-esophageal sample and the rumen fractions samples from fistula.

### Rumen metabolome

A total of 185 known and 236 unknown primary metabolites were identified in the rumen content. The concentration of metabolites discriminating the sampling methods among rumen content samples was used for the partial least squares discriminant analysis (PLS-DA, Fig. 4). Overall, PLS-DA with known metabolites indicate the metabolome composition and concentration of samples collected through the oro-esophageal procedure and the combined fluid-particulate fractions are similar. When comparing the fluid versus the particulate fractions separately, PLS-DA with known metabolites indicate the metabolome composition of these two fractions are considerably different. The PLS-DA for unknown metabolites illustrated a moderate overlap of oro-esophageal samples with combined fluid-particulate contents. The same pattern of distinction is observed for PLS-DA with unknown metabolites when comparing the fluid and particulate fractions separately as reported for known metabolites. An interesting pattern that is observed in both PLS-DA with known and unknown metabolites is that despite the similarity of metabolome composition between samples collected through the oro-esophageal procedures and the rumen fistula fluid-particulate fractions combined is the dispersion of the data. The PLS-DA shows the oro-esophageal procedure dispersion is greater than the rumen fistula fluid-particulate fractions combined, which indicated the former may require a greater number of experimental units to have a representative overview of the population compared to the latter.

**Figure 4.**
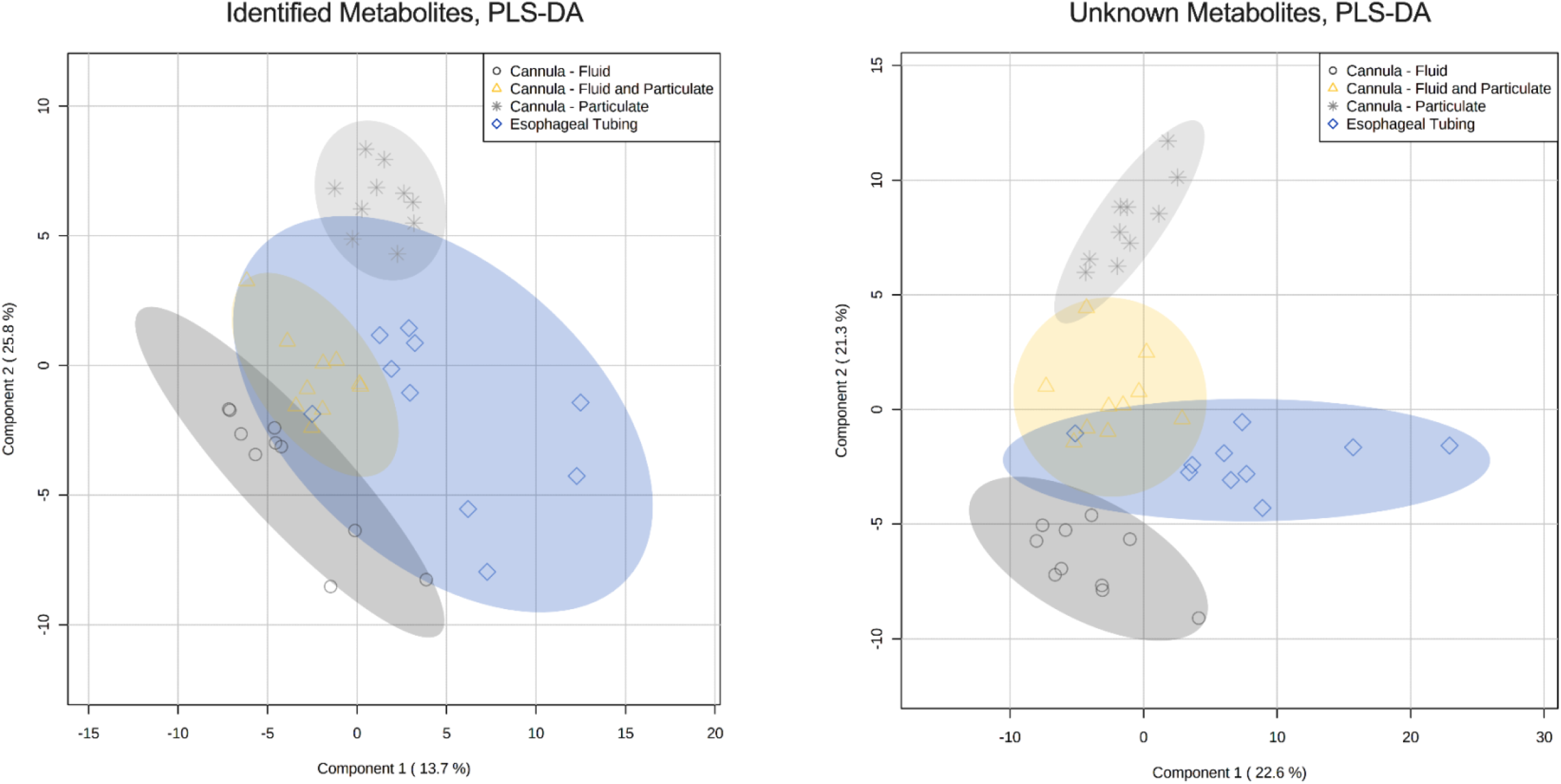
Partial least square-discriminant analysis (PLS-DA) of the ruminal metabolites from ruminal samples collected through an oro-esophageal tubing, and the fractions fluid, particulate, and combined fluid-particulate from rumen fistula.

**Figure 5.**
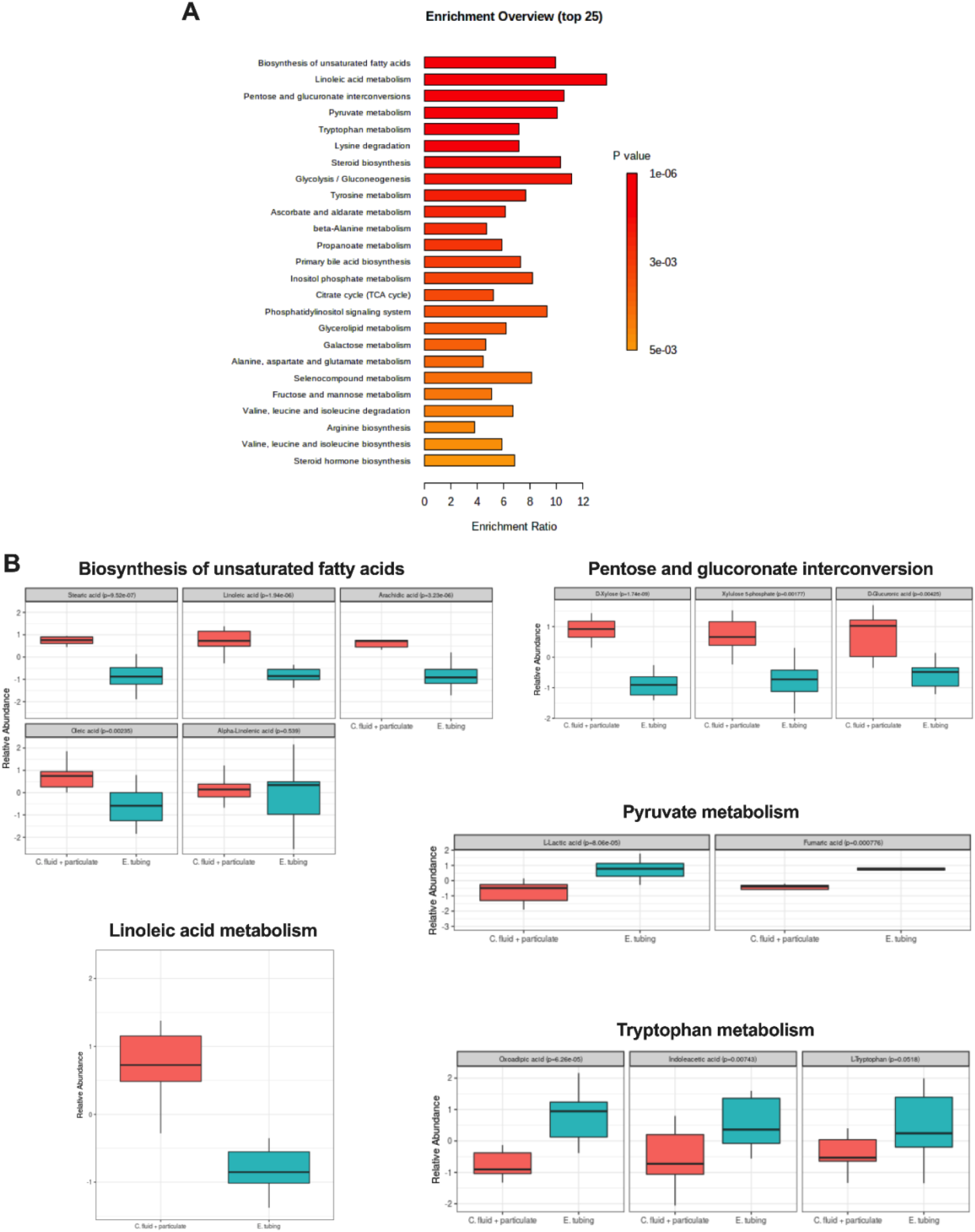
Hierarchical of ruminal metabolomic by enrichment pathways overview (top 25) (A) and Metabolites (B) from ruminal samples collected through an oro-esophageal tubing and the combined fluid-particulate from rumen fistula only as a direct comparison between a newer rumen sampling methodology with the traditional gold standard used in ruminant studies. Statistical differences were declared at *P* ≤ 0.05.

Clustering the metabolome composition and concentration together into known metabolic pathways allowed the understanding of whether any of the sampling methods would affect or not the study of major topics of nutrient studies in ruminants. In the current study, differences between sampling methods were found to mainly affect metabolites associated with the biosynthesis of unsaturated fatty acids (Fig. 5A). The concentrations of stearic, linoleic, arachidic, and oleic acids were greater in the rumen fistula fluid-particulate fractions combined than in the oro-esophageal samples (Fig. 5B). The pathway associated with linoleic acid metabolism was the second most enriched pathway followed by the pentose and glucuronate interconversions (Fig. 5B), where the concentration of D-xylose, xylulose 5-phosphate, and D-glucoronic acid were greater in the rumen fistula fluid-particulate samples than oro-esophageal ones. Following the pathways of pyruvate and tryptophan metabolism with the concentration of oxoadipic acid, indoleacetic acid, L-tryptophan metabolites greater in oro-esophageal samples than in the rumen fistula fluid-particulate ones.

## DISCUSSION

The current study sheds light on the impact of different sampling methods to characterize the rumen microbiome and metabolome concurrently in dairy cows. Here, the microbiome findings suggest that studies designed to use the oro-esophageal probe to collect rumen samples for microbiome evaluation can yield similar results for different fractions of rumen contents collected through the rumen fistula. Despite the similarities for the rumen microbiome, these data also suggest that the overall rumen metabolome of the oro-esophageal procedure can be represented by the combined fluid-particulate fractions collected from rumen fistula, but that the metabolome composition of the independent fractions from fistula is distinct. These findings highlight that methodologies and which rumen fractions to be used are important considerations when studying specific metabolites. Nonetheless, the microbiome and metabolome of oro-esophageal and rumen fistula fluid-particulate combined samples were generally similar, highlighting once again the interchangeable similarity between both methods when evaluating ruminal microbial and metabolic changes.

As expected and previously demonstrated (29), the pH of rumen samples was greater in the oro-esophageal tubing than in contents from the rumen fistula. The findings of the current study confirm that samples collected with an oro-esophageal tubing may, in fact, present a higher pH. Still, despite this difference, the overall microbiome for the oro-esophageal and rumen fistula fluid-particulate combined samples did not diverge. Therefore, if the goal of researchers is simply to characterize the rumen microbiome changes using 16S rDNA in a single time in relation to feeding, the current findings suggest that the variation in pH has a negligible impact on the characterization of the rumen microbiome present in fluid and particulate fractions of the rumen.

The effect of rumen fractions and the methodologies were not significant for most bacterial communities. Even the richness of particulate and fluid-particulate samples containing degraded fiber when compared to fluid did not differ from rumen contents and between sampling methods as described before (16, 30). These results are in agreement with other studies that compared these methodologies to collect rumen samples in different hours after feeding (30), the collection in different sites (16), and in pre-weaned calves (31). However, Deusch et al. 2017 (32) found a significance change on bacterial community arrangement over different rumen fractions and diets, which is probably more likely associated with the different diets than sites.

The most prevalent phyla were consistently Bacteroidetes followed by Firmicutes in oro-esophageal and rumen contents collected using the fistula (16, 32). In a study comparing the rumen microbiome from 32 different ruminant species in 35 different countries all over the world (33), these two phyla were the most abundant and part of a core microbiome considering variations in diet and host. Specifically evaluating sampling methodologies, De Assis and collaborators (2020) (30) found an increase in the relative abundance of Firmicutes and a decrease of Bacteroidetes in samples from rumen fistula when compared to stomach tubing over time. The differences in the study from De Assis and collaborators (30) were reported to be associated with the time of collection (no differences up to 4 hours) and pH due to potential contamination with saliva. In the current study, the lack of difference was noted beyond 4 hours in a single time point collection. Thus, precautions to avoid repeated interaction with rumen contents and exposure to oxygen may mitigate potential causes of differences between the two techniques. Also, in the current study, the Bacteroidetes to Firmicutes ratio was not significant among sampling techniques, indicating such variation may not happen in a more controlled setting. At the genus level, Prevotella was the most abundant genus found in the rumen samples of the current study, which has been widely reported in other rumen studies as well (8, 9, 16), including rumen microbiome studies with a large number of samples analyzed (33). Prevotella is one of the major microbes responsible for the degradation of starch and protein and plays an essential role in volatile fatty acids biosynthesis. This genus is also associated with cows with high milk yield and milk protein content which were the characteristics of the cows used in our study (14).

Another difference observed by De Assis and collaborators (2020) (30) was regarding the variation in beta-diversity in samples from the oro-esophageal technique and those collected using rumen fistula, showing a larger variation in the former. This larger variation was not found in the microbiome analysis of our study, but the metabolome one had a similar larger variation that is described later in this discussion. Henderson et al. (2013) (34) comparing different methodologies for the extraction and sequencing of rumen bacteria and archaea communities show that there is a variation depending on the protocol to be used, but that this variation is not large enough to be present in dimension reduction analysis such as the principal coordinate ones (PCoA) reported here. In both cases, even considering such variation in previous microbiome studies, the centroid of populations are similar and often overlap in these analyses, meaning differences between the oro-esophageal technique and the whole sample from rumen fistula considering a reasonable sample size may not be as large as previously reported. Other factors that could potentially change ruminal parameters and need careful consideration when using the oro-esophageal technique are the depth of the inserting tube and the probe size used to collect the rumen content (34, 35). The probe used in the current study had openings large enough to pass particulate fractions that could account for microbial populations attached to feedstuff, which overall may have contributed to such small differences in microbial populations from this study.

Regarding metabolite dispersion across different techniques and types of samples, the current study revealed that liquid and particulate are distinct while the liquid-particulate and oro-esophageal are similar, as shown for the microbiome in a previous study (30). Therefore, even though variation in sample composition exists, their centroids do not differ, suggesting that larger sample size studies, which is a reason and potential advantage of using the oro-esophageal technique, may help reduce the misrepresentation of populations. For specific metabolites, large differences were observed mainly between the fluid and particulate fractions of the rumen content collected through rumen fistula, possibly due to the nature of fractions and nutrients that generate their respective end-products of fermentation. However, when considering samples that contains similar physical composition such as the oro-esophageal and fluid-particulate combined from rumen fistula, the difference is mostly associated with an overall contribution of small metabolite variations, which changed some metabolic pathways as reported here.

Sampling through the fistula has been traditionally used because it allows the collection of a more representative sample of the contents. In the case of the oro-esophageal tubing that is not filtered, the opening in the collection probe also allows the collection of feed particles in such samples. This might explain why the rumen metabolite composition from the oro-esophageal technique was similar to the fluid-particulate samples collected by rumen fistula. However, because the sample is not taken as a mixed one from all sites of the rumen but at random, a slight variation could be introduced, and more samples may be necessary to characterize metabolome phenotypes more accurately. Thus, due to the close relationship between ruminal metabolites with different pathways and even their direct presence in different ones (36), the current study suggests rumen metabolome studies should be carefully designed, and an adequate number of experimental units can be a factor to be considered to avoid such problems.

An example is unsaturated fatty acids, which are synthesized by aerobic and anaerobic mechanisms depending on the organism (37). Not only the oxygen but the environment, temperature, and nutrition can modify the composition of the lipid molecule (38). These factors can explain the greater concentrations of the metabolites (stearic, linoleic, arachidic, oleic and alpha-linolenic acids) in combined fluid-particulate samples than in oro-esophageal samples. Changes in the rumen environment due to the oxygen circulation through the fistula for this specific pathway may alter lipid metabolism pathways. If the ultimate goal of a study is the characterization of specific lipids, the oro-esophageal technique may be an advantageous approach. The difference in unsaturated fatty acids exemplifies how these changes in metabolome composition are less likely to follow the same pattern as those found in the microbiome. There are also metabolites derived from other microorganisms, different plant materials, or even the host (11), which could potentially change the study’s outcome, but the contribution of these factors was beyond the scope of the current study.

### Conclusion remarks

In conclusion, the current study indicates that despite having greater rumen content pH, the oro-esophageal procedure did not present major microbiome or metabolome composition differences when compared to the combined fluid-particulate samples from rumen fistula. Most differences were observed when comparing metabolites from the fluid and particulate fractions separately. Lastly, small variations in some rumen metabolites may potentially change specific metabolic pathways in the rumen. Thus, studies looking at specific ruminal pathways associated with the rumen microbiome should carefully consider the sampling method to be used.

## ACKNOWLEDGMENTS

We acknowledge Sara Knollinger for the available data and the cows, Ahmer A. Elolimy for helping us during the collections, Mohamed M. Zeineldin for the help with the bead beater machine, and Phillip Peixoto for the abstract poster presentation at the ADSA Meeting in 2019.

